# TimiGP-Response: the pan-cancer immune landscape associated with response to immunotherapy

**DOI:** 10.1101/2024.06.21.600089

**Authors:** Chenyang Li, Wei Hong, Alexandre Reuben, Linghua Wang, Anirban Maitra, Jianjun Zhang, Chao Cheng

## Abstract

Accumulating evidence suggests that the tumor immune microenvironment (TIME) significantly influences the response to immunotherapy, yet this complex relationship remains elusive. To address this issue, we developed TimiGP-Response (TIME Illustration based on Gene Pairing designed for immunotherapy Response), a computational framework leveraging single-cell and bulk transcriptomic data, along with response information, to construct cell-cell interaction networks associated with responders and estimate the role of immune cells in treatment response. This framework was showcased in triple-negative breast cancer treated with immune checkpoint inhibitors targeting the PD-1:PD-L1 interaction, and orthogonally validated with imaging mass cytometry. As a result, we identified CD8+ GZMB+ T cells associated with responders and its interaction with regulatory T cells emerged as a potential feature for selecting patients who may benefit from these therapies. Subsequently, we analyzed 3,410 patients with seven cancer types (melanoma, non-small cell lung cancer, renal cell carcinoma, metastatic urothelial carcinoma, hepatocellular carcinoma, breast cancer, and esophageal cancer) treated with various immunotherapies and combination therapies, as well as several chemo- and targeted therapies as controls. Using TimiGP-Response, we depicted the pan-cancer immune landscape associated with immunotherapy response at different resolutions. At the TIME level, CD8 T cells and CD4 memory T cells were associated with responders, while anti-inflammatory (M2) macrophages and mast cells were linked to non-responders across most cancer types and datasets. Given that T cells are the primary targets of these immunotherapies and our TIME analysis highlights their importance in response to treatment, we portrayed the pan-caner landscape on 40 T cell subtypes. Notably, CD8+ and CD4+ GZMK+ effector memory T cells emerged as crucial across all cancer types and treatments, while IL-17-producing CD8+ T cells were top candidates associated with immunotherapy non-responders. In summary, this study provides a computational method to study the association between TIME and response across the pan-cancer immune landscape, offering resources and insights into immune cell interactions and their impact on treatment efficacy.

## Main

In the past decade, immunotherapy, as exampled by immune checkpoint inhibitors (ICIs), has revolutionized treatment across different cancers^1–3^. However, only a small fraction of patients can achieve long-term clinical benefits, while the majority of patients have limited benefit^4,5^. Overall, the response rate to ICI monotherapy in unselected populations is only about 15 to 20 percent^5^. Consequently, substantial efforts have been made to enhance comprehension of underlying mechanisms associated with immunotherapy resistance.

Accumulating evidence suggests that the tumor immune microenvironment (TIME) significantly influences the benefit from immunotherapy, yet this complex relationship remains elusive^6–8^. The roles of various immune cells, their subtypes, and their interactions in the response to immunotherapy add layers of complexity that are not yet fully understood. To more comprehensively study the TIME, traditional methods like immunohistochemistry (IHC), flow cytometry, and emerging techniques such as imaging mass cytometry (IMC) are limited in the markers and cell types analyzed simultaneously. On the other hand, transcriptomic profiling techniques offer a promising approach with broader coverage and more flexible analysis. Single-cell RNA sequencing (scRNA-seq) provides unparalleled resolution, enabling the dissection of cellular diversity at a high level, but its clinical application is hindered by prohibitive costs and stringent sample quality requirements, often resulting in sequencing small cohorts of specimens^9,10^. In contrast, bulk transcriptomic profiling, with its established methodologies and clinical platforms, presents a cost-effective approach capable of accommodating larger cohorts, albeit at the sacrifice of resolution^11–13^. In light of these factors, it becomes evident that effectively leveraging the high resolution of single-cell RNA sequencing and the larger cohort capacities of bulk transcriptomic profiling could enhance our ability to comprehensively explore the association between the TIME and immunotherapy response. This necessitates computational methods that integrate the strengths of both techniques.

To fill this void, we developed TimiGP (<u>T</u>umor Immune <u>M</u>icroenvironment Illustration based on <u>G</u>ene <u>P</u>airs), a computational framework under the hypothesis that the dynamic equilibrium of the immune system plays a pivotal role in determining the clinical outcomes of patients^14,15^. Using TimiGP, we could assess cell-cell interactions and the prognostic value of immune cells through survival statistics, bulk transcriptomic data, and cell-type markers derived from the scRNA-seq analysis. In our previous research, we demonstrated its capability to uncover immune cell prognostic associations and interactions at different resolutions and across various cancer types.

Moreover, the TimiGP framework has been demonstrated as a methodology adept at leveraging the strengths of both scRNA-seq and bulk transcriptomic profiling^14,15^. Considering the rationale behind TimiGP is beyond solely prognosis^14,15^, in this study, we extended the TimiGP framework to explore the association between the TIME and the response to immunotherapy (TimiGP-Response, Fig. 1a, Supplementary Fig. 1, See Methods for more details).

**Fig. 1.**
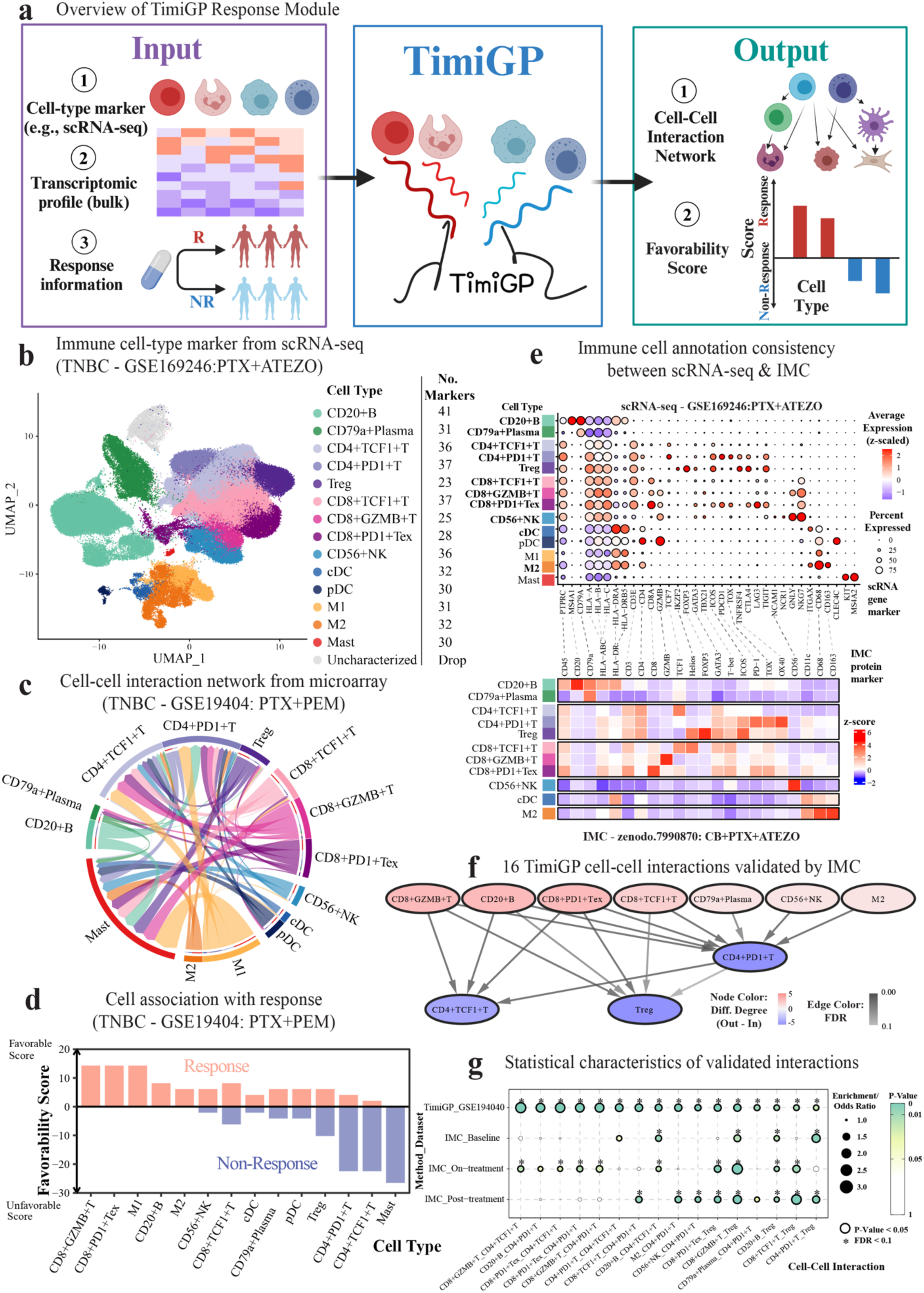
Exemplary applications and independent validation of TimiGP-Response in TNBC. a. Schematic depicting the TimiGP - Response. This figure was created with BioRender.com. b. UMAP plot of 14 immune cell types tailored for TNBC using scRNA-seq. c-d. TIME associated with anti-PD-1 response depicted by TimiGP – Response Module. c. Chord diagram of all cell-cell interactions. The arrow (X → Y) represents cell-cell interactions from favorable cell X (color of the outer ring and arrow) to unfavorable cell Y (color of the inner ring), indicating a high X/Y ratio is associated with a favorable response. The wider the arrow is, the smaller the FDR is. d. Bar plot of the favorability score to prioritize cell type associated with response (orange) or non-response (blue). e. The similar expression profile between scRNA-seq and IMC defined cell types. The upper dotplot displays the gene expression profile of 14 cell types identified in the scRNA-seq data. On the x-axis, cell type gene markers are listed, while the y-axis represents each cell type. The expression values are z-scaled and depicted as colors, with the size of the dots representing the percentage of marker expression in each cell type. Cell types also identified in the IMC data are highlighted in bold. The bottom heatmap exhibits the z-scored protein expression, with the y-axis representing cell types and the x-axis representing protein markers. Dotted lines connect the protein names with their corresponding gene names. f-g. 16 TimiGP cell-cell interactions validated by independent IMC data. f. The cell-cell interaction network representing 16 interactions identified by TimiGP-Response and validated by IMC data. The node represents the cell type, and its color shows the difference between out-degree and in-degree. The edge represents cell-cell interaction, and its transparency shows the FDR calculated from TimiGP - Response. g. Statistical properties of those16 cell-cell interactions. The x-axis represents cell interactions, while the y-axis displays TimiGP – Response and IMC results obtained from different time points. The TimiGP results consist of the enrichment ratio and corresponding p-value, derived from the TimiGP framework via enrichment analysis. The IMC results include the odds ratio and corresponding p-value, obtained through logistic regression, where each cell ratio calculated based on the image serves as the independent variable and the therapy response as the dependent variable. Abbreviation of cell types: B, B cell; Plasma, Plasma cell; T, T cell; Treg, Regulatory T cell; Tex, Exhausted T cell; NK, Natural killer cell; DC, Dendritic cell; cDC, Conventional DC; pDC, Plasmacytoid dendritic cell; M0, Non-activated macrophage; M1, Pro-inflammatory macrophage; M2, Anti-inflammatory macrophages; Mast, Mast cell. Abbreviation of treatment: PTX, paclitaxel (chemotherapy); PEM, pembrolizumab (anti-PD-1 immunotherapy); CB, Carboplatin(chemotherapy); ATEZO, atezolizumab (anti-PD-L1 immunotherapy).

Considering the multi-omics data availability, our study began on cohorts of triple-negative breast cancer (TNBC) and immune checkpoint inhibitors targeting the interaction between Programmed Cell Death Protein 1 (PD-1) and Programmed Death-Ligand 1 (PD-L1). In order to examine the TimiGP - Response, We collected data from several TNBC cohorts, including scRNA-seq^16^, bulk transcriptomic profiling^17^, IMC^18^, and the corresponding response information related to treatments combining anti-PD-1/PD-L1 therapies with chemotherapy.

We first reanalyzed publicly available scRNA-seq data^16^ and identified cell-type markers representing 14 distinct immune cell populations (Fig. 1b, Supplementary Fig. 2-3, Supplementary Table 1). Next, we conducted a comparative analysis of the expression profiles across different cell types and subsequently generated marker annotations for these cell types (See Methods for more details). This cell-type annotation customized for the TNBC and anti-PD-1/PD-L1 treatment was subsequently fed into the TimiGP-Response as the first input to analyze the second input, microarray profiling of pre-treatment samples from a larger TNBC cohorts^17^, together with the corresponding treatment response, the third input.

As a result, TimiGP generated a comprehensive cell-cell interaction network, where each arrow indicated an interaction from a favorable to an unfavorable cell type (Fig. 1c, Supplementary Table 2). For instance, the interaction from CD8+ GZMB+ T cells to Tregs suggested a high ratio between these cell types associated with treatment responders. Employing network analysis, TimiGP computed favorability scores, prioritizing cell types associated with either treatment response or non-response (Fig. 1d). Among those cell types, we found CD8+ GZMB+ T cells and CD8+ PD1+ exhausted T cells (Tex) potentially correlated with treatment responders, aligning with the therapeutic rationale targeting PD-1 and reversing T cell exhaustion^1^.

To orthogonally validate the findings from the TimiGP – Response, we reanalyzed a publicly available immune profiling dataset by IMC from an independent TNBC cohort^18^. We identified 11 immune cell types exhibiting expression patterns analogous to those defined in the scRNA-seq study (Fig. 1e). To validate the relationship between cell types and immunotherapy responses in TNBC, computed associations between individual cell densities or the ratios among 11 different immune cell types and treatment response in this IMC dataset (Supplementary Table 3). Subsequently, we compared the IMC findings with our TimiGP-inferred cell-cell interactions derived from microarray data. As expected, CD8+ GZMB+ T cells was also significantly associated with responders in pre-treatment samples from the IMC cohort (Supplementary Fig. 4). Of note, 16 TimiGP cell-cell interactions were validated using this independent IMC dataset, thereby supporting the method’s reliability. Moreover, these cell-cell interactions, such as CD8+ GZMB+ T cells → Treg, CD20+B cells → Treg and CD8+ PD1+ Tex → Treg interactions, may hold promise as cell ratio-based features for patient selection and monitoring treatment efficacy during immunotherapy (Fig. 1g).

Following the validation of TimiGP-Response in TNBC, we employed this methodology to illustrate the pan-cancer TIME landscape in relation to immunotherapy response. We collected public datasets including over 3,000 patients spanning 7 distinct cancer types and subjected to different immunotherapies (Supplementary Table 4). We adhered to the criteria outlined in the original study to define the response and analyzed them separately with TimiGP-Response at different resolution (Fig. 2a).

**Fig. 2.**
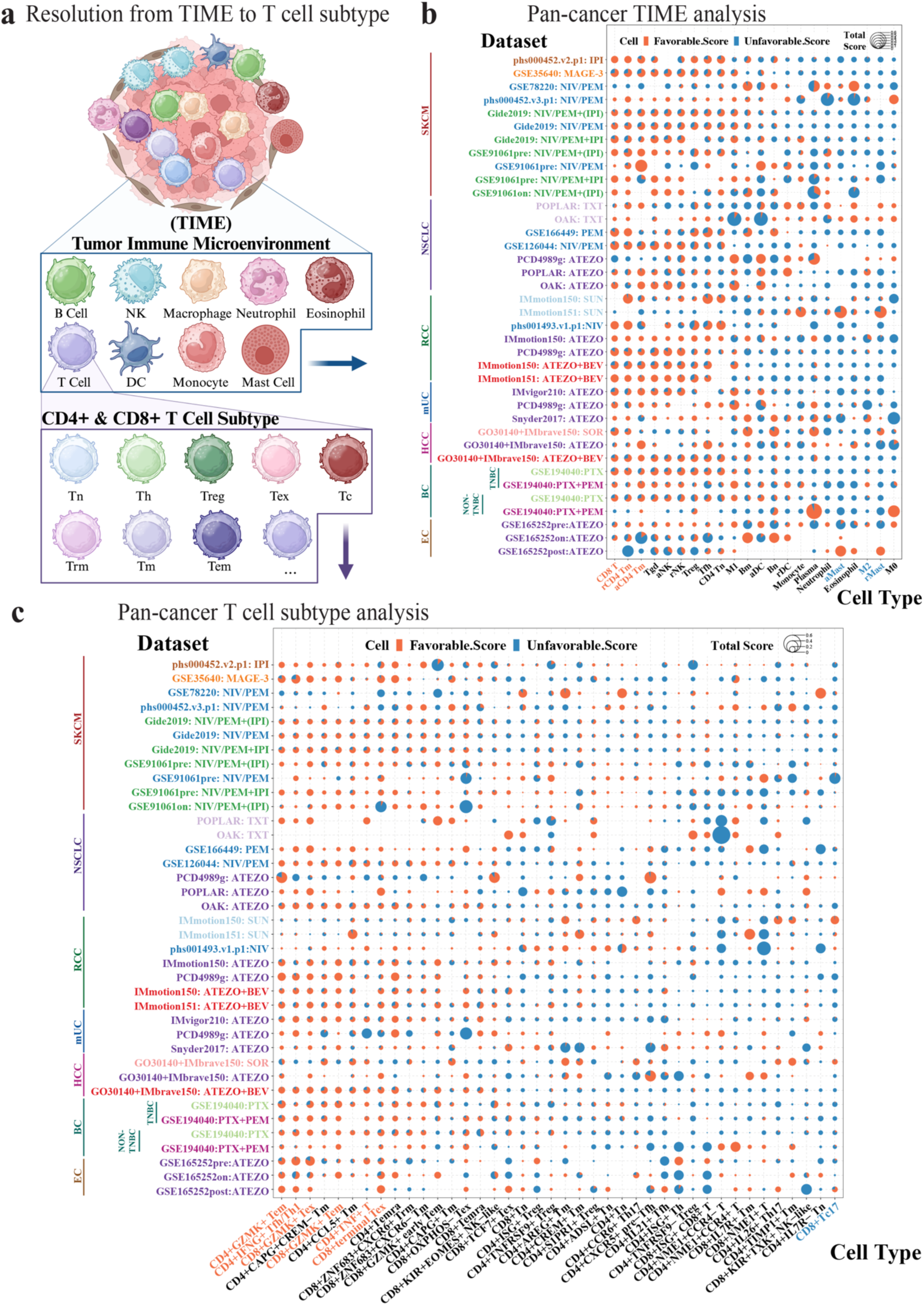
Pan-cancer TIME and T cell subtype landscape associated with response to immunotherapy. a. Schematic demonstrating the resolution of pan-cancer analysis from the TIME to T cell subtypes. This figure was created with BioRender.com. b-c. TimiGP-Response illustrates pan-cancer (b) TIME and (c) T cell subtypes landscape associated with immunotherapy response. The scatter pie chart displays the favorability score exported by TimiGP - Response Module, which estimates the association of immune cell types defined by (b) modified LM22 annotation or (c) T cell subtypes (x-axis) with immunotherapy responders (favorable score, orange) and non-responders (unfavorable score, blue). The cell types discussed in the results were highlighted with orange and blue colors. There are 7 cancer types in this analysis: SKCM (melanoma), NSCLC (non-small cell lung cancer), RCC(renal cell carcinoma), mUC (metastatic urothelial cancer), HCC (hepatocellular carcinoma), BC (breast cancer; TNBC, triple-negative breast cancer), EC (esophageal cancer). The datasets (y-axis) are labeled in the format “dataset ID: drug”, with immunotherapy datasets shown in dark colors, and chemotherapy and targeted therapy datasets shown in light colors. The immunotherapy includes anti-CTLA4 (IPI: ipilimumab), MAGE-A3 vaccine, anti-PD-1 (NIV: nivolumab, PEM: pembrolizumab), combination of anti-CTLA4 and anti-PD-1, anti-PD-L1 (ATEZO: atezolizumab), and combination of anti-PD-L1 and anti-VEGF (BEV: bevacizumab). The chemotherapy includes docetaxel (TXT) and paclitaxel (PTX). The targeted monotherapy includes sunitinib (SUN) and sorafenib (SOR) which target receptor tyrosine kinases (RTKs). Abbreviation of cell types: Bn, Naïve B cell; Bm, Memory B cell; Plasma, Plasma cell; T, T cell; Tn, Naïve T cell; Th, Helper T cell; Th1, Typer Th; Th17, Type 17 Th; Tfh, Follicular Th; Treg, Regulatory T cell; Tex, Exhausted T cell; Tc17, Type 17 T cell; Tm, Memory T cell; Tem, Effector memory T cell; Trm, Tissue-resident memory T cell; Temra, Terminally differentiated effector memory or effector T cell; Tgd, Gamma delta T cell(Tγδ); NK, Natural killer cell; DC, Dendritic cell; M0, Non-activated macrophage; M1, Pro-inflammatory macrophage; M2, Anti-inflammatory macrophages; Mast, Mast cell; r, Resting; a, Activated.

First, we utilized cell-type markers adapted from Bindea et al.^19^, encompassing two critical cell types: cytotoxic cells, serving as positive controls indicative of anti-tumor activity, and tumor cells, designated as negative controls associated with non-responders^14,15^. By integrating these control cell types into our analysis, we aimed to assess the performance of TimiGP-Response in a pan-cancer context.

As shown in Supplementary Fig. 5, T cells and cytotoxic cells demonstrated a consistent association with immunotherapy responders, while tumor cells were predominately associated with non-responders across nearly all datasets. These results align with the designed controls and the rationale behind immunotherapy targeting T cell responses, indicating the robustness of our approach in discerning immune profiles associated with treatment outcomes across various cancers.

Given the potential bias arising from assigning cytotoxic markers of CD8 T cells and NK cells to the specific cytotoxic cell, we subsequently portray the TIME utilizing the modified LM22 signature^14,15,20^. This signature includes activating and resting immune cell states and has undergone extensive validation^20,21^. As shown in Supplementary Fig. 6, we observed significant similarities in cell-cell interaction networks among immunotherapy cohorts within the same cancer types, regardless of treatment regimens (e.g., MAGE-3 vaccine, anti-CTLA-4 and anti-PD-1 treatment in SKCM). Furthermore, this similarity extended across distinct cancer types that underwent similar regimens (e.g. SKCM, RCC, NSCLC treated with anti-PD-1 therapy). Of note, while immunotherapy cohorts exhibited substantial similarity, chemotherapy, and targeted therapy cohorts displayed pronounced differences from immunotherapy cohorts. These findings collectively support the robustness of our methodology, agnostic to tumor types but specific to immunotherapy, in line with the design of TimiGP-Response with the focus on immune cells.

Considering the similarity observed in the cell-cell interaction network across immunotherapy datasets, our next step is to leverage the TimiGP-Response to discern potential pan-cancer features indicative of response to the immunotherapy (Fig. 2b). Across the seven cancer types and various immunotherapies, the major immunotherapy target, CD8 T cells, along with resting and activated CD4 memory T cells, are consistently associated with responders (Fig. 2b). This is consistent with the central role of CD8 T cells in tumor surveillance and cytotoxicity^22^, as well as the immunomodulatory functions of CD4 memory T cells in enhancing immune responses and serving as a reservoir for future activation upon encountering antigens^23^.

Conversely, anti-inflammatory (M2) macrophages and resting mast cells are associated with non-responders to immunotherapy (Fig 2), which aligns with previous reports that M2 macrophages exert immunosuppressive effects, hindering the effectiveness of immunotherapy and fostering tumor immune evasion and progression^24–26^. As regards to mast cells, tumor-infiltrating mast cells have been linked to resistance against anti-PD-1 therapy in mouse models ^27–29^.

Hereto, we applied the TimiGP-Response to illustrate the association of the pan-cancer TIME landscape with immunotherapy response. Our results not only demonstrate the superior performance of our methodology in pan-cancer analysis but also provide comprehensive insights into the association between distinct innate and adaptive immune cell states and responses to different immunotherapy across seven cancer types.

Given that the main targets of the immunotherapy in these datasets are T cells, and our analysis at the TIME resolution also highlights the importance of T cells in response to immunotherapy, we next focus on T cells for a higher resolution, including 40 T cell subtypes as defined in the scRNA-seq study^30^.

As shown in Fig. 2c, CD8+GZMK+ exhausted T cells (Tex) and CD8+terminal Tex emerged as pivotal cell types associated with immunotherapy responders across nearly all cancer types and various immunotherapy regimens. These cells play a critical role in directly eliminating tumor cells but can succumb to exhaustion, characterized by decreased effector function. They can be reversed by immunotherapies, especially ICIs, to unleash their full potential against cancerous cells^1,31,32^. This finding is consistent with our previous analysis, which not only aligns with the rationale behind immunotherapy but also demonstrates the reliability of our method.

CD8+ and CD4+ GZMK+ effector memory T cells (Tem) were also identified as associated with immunotherapy responders. GZMK (encoding Granzyme K) serves as a marker indicative of anti-tumor cytotoxic effector function. These cytotoxic effector memory T cells have been reported to play a crucial role and are positively linked to immunotherapy response^22,33,34^.

In addition, CD4+IFNG+ follicular/type 1 dual helper T cells (Tfh/Th1) and CD4+TNF+ T cells were identified to demonstrate a favorable correlation with immunotherapy response. IFNG and TNF are key cytokines influencing tumor cell viability and regulating anti-tumor immune responses, potentially enhancing the effectiveness of cancer immunotherapy^23,35–39^.

On the other hand, for immunotherapy non-responders, CD8+Tc17 (IL-17 producing CD8+ T cells) emerges as a top candidate. Although still under debate, CD8+ Tc17 has been reported to promote tumor progression^40,41^, and contribute to CD8+ T cell exhaustion^42,43^. Blocking CD8+ Tc17 may decrease CD8+ T cell exhaustion, inhibit tumor growth and enhance response to anti-PD-1 immunotherapy in mouse models^42,44,45^. Our results consistently demonstrate its negative association with immunotherapy response highlighting the promise of TimiGP-Response in identifying cellular features associated with immunotherapy failure.

Taken together, our findings further delineate the T cell landscape associated with immunotherapy response, providing insights for developing novel biomarkers to select patients who may benefit from immunotherapy as well as identifying therapeutic targets to overcome resistance to immunotherapy.

## Discussion

In this study, we expanded the TimiGP framework beyond prognosis to encompass immunotherapy response, culminating in the TimGP-Response Module. This module integrates scRNA-seq, bulk transcriptomic data, and response information, facilitating the inference of cell-cell interaction networks and the identification of priority cell types linked to responders and non-responders to immunotherapy. By leveraging the high-resolution capabilities of scRNA-seq and compensating for the limitations of small cohort sizes with bulk data, the TimGP-Response Module effectively bridges the gap in existing methodologies by accommodating multi-modality data, including multi-omics and clinical information.

Notably, despite variations in different cohorts and profiling technologies, we successfully validated 16 cell-cell interactions identified from microarray using TimiGP-Response in independent cohorts with IMC data. This validation suggests that our statistical cell-cell interactions may also exhibit spatial relationships, thereby indicating the efficacy of our method. Our pan-cancer analysis, which robustly identifies the positive control, cytotoxic cells, associated with immunotherapy responders and the negative control, tumor cells, associated with non-responders, also supports the performance of this method.

Our comprehensive analysis involved the collection of data from over 3000 patients spanning 7 cancer types, with a primary emphasis on immunotherapy. Through profiling the immune landscape associated with immunotherapy response across various resolutions, we have generated a valuable resource comprising both curated datasets and resulting cell-cell interaction networks within TIME and T cell subtypes. This resource is poised to benefit the research community by enhancing our understanding of the mechanisms underlying immunotherapy resistance.

Moreover, we highlighted several immune cell types strongly correlated with responders and non-responders, which aligns with established knowledge (e.g., CD8+ GZMK+ Tex) or previous experimental findings (e.g., CD8+ Tc17). Although further experimental validation is warranted to solidify these intriguing findings, the consistent identification of cell types across diverse cancer types and immunotherapy regimens increases our confidence. Our method potentially contribute to select biomarkers, which may benefit patient stratification to optimize immunotherapy outcomes and to identify novel therapeutic targets for non-responders. Additionally, other cell types (e.g., CD4+ISG+ Treg), albeit not described in detail, may also play important roles in immunotherapy response across various cancers or exhibit distinct behaviors in specific cancer types or treatments (e.g., CD4+ISG+ Treg), which are worth further experimental investigation.

While our primary focus lies in elucidating the association between immune cells and immunotherapy response, the adaptability of our method allows for the exploration of other cell types within the tumor microenvironment and their relationships with alternative therapies such as chemotherapy, radiotherapy, and targeted therapy through the customization of cell types of interest using scRNA-seq data. We anticipate that our method will serve as a valuable tool for unraveling the complex interplay between the tumor microenvironment and all types of treatment response, ultimately advancing precision medicine.

## Supplementary information

Supplementary Figure 1 | The rationale and framework of TimiGP-Response adapted from TimiGP-Prognosis. Related to Figure 1

Supplementary Figure 2 | Overview of clinical features and cell proportions reanalyzed from GSE169246. Related to Figure 1

Supplementary Figure 3 | The expression of cell-type markers in UMAP. Related to Figure 1

Supplementary Figure 4 | The association between immune cell density and immunotherapy response

Supplementary Figure 5 | Robust performance of TimiGP-Response in pan-cancer analysis. Related to Figure 2

Supplementary Figure 6 | High similarities of TIME cell-cell interaction networks across cancer types and treatments. Related to Figure

Supplementary Table 1 | TNBC Cell-type markers tailored with single-cell RNA-seq

Supplementary Table 2 | Statistical details of cell-cell interactions exported by TimiGP-Response

Supplementary Table 3 | Statistical details of cell-cell interactions analyzed using IMC data

Supplementary Table 4 | Pan-cancer immunotherapy datasets

## Methods

### Data collection and preprocessing

#### Triple negative breast cancer (TNBC) multi-omics datasets

In the case of TNBC, we gathered single-cell RNA-seq (scRNA-seq) data^16^, microarray data^17^, and imaging mass cytometry (IMC) data^18^ from different TNBC cohorts that received similar treatments.

The preprocessed scRNA-seq data (GSE169246) was obtained from the Gene Expression Omnibus (GEO)^16^. This dataset consists of 14 patients with advanced TNBC who were treated with either paclitaxel alone or in combination with atezolizumab (anti-PD-L1 therapy). It is utilized to tailor cell-type markers for TimiGP-Response, specifically for TNBC, and for immunotherapy targeting PD-1 and PD-L1 interactions in conjunction with chemotherapy. In the dataset, 7 patients received the combination of chemotherapy and immunotherapy. Of these, 6 have both pre- and post-treatment samples, while 1 has only pre-treatment samples. Additionally, 7 patients received chemotherapy alone, with 6 having both pre- and post-treatment samples and 1 having only post-treatment samples.

The preprocessed microarray data (GSE194040) was downloaded from the GEO. For the TimiGP-Response analysis tailored to TNBC, we used data from 26 TNBC patients treated with a combination of paclitaxel and pembrolizumab (anti-PD-1)25, along with their corresponding treatment responses^17^. The TNBC cohorts was characterized by the absence of estrogen receptor (ER), progesterone receptor (PR), and human epidermal growth factor receptor 2 (HER2) expression^46,47^. Thus, we filtered the patients to include only those who were HER2 negative (HER2 = 0) and hormone receptor (ER and PR) negative (HR = 0). Based on the pathological complete response (pCR) status provided by the original study, we categorized patients with a complete response (pCR = 1) as responders (N=13) and those with a failed complete response (pCR = 0) as non-responders (N=13).

The preprocessed IMC data were requested from via Zenodo data repository (https://doi.org/10.5281/zenodo.7990870). To orthogonally validate the results from TimiGP-Response, we selected 127 TNBC patients treated with chemotherapy plus anti-PD-L1 immunotherapy (carboplatin, nab-paclitaxel and atezolizumab)^18^. The samples were obtained from FFPE tissues at different time points: baseline (N = 113, 56 responders and 57 non-responders), on-treatment (N = 97, 48 responders and 49 non-responders), and post-treatment (N = 104, 56 responders and 48 non-responders). Among them, 73 patients had the samples from all three time points. Based on response information provided by the original study, we categorized patients with pCR as responders and those with residual disease (RD) as non-responders.

#### Pan-cancer bulk transcriptomics datasets

After examining the performance of TimiGP-Response in TNBC, we extended our analysis to a pan-cancer setting. We gathered transcriptomic profiles from publicly available resources including 3,410 cancer patients, along with treatment response data. The sources of those datasets were listed in Supplementary Table 4 (columns “NCT”, “DOI”, “Authors”, “DatabaseID”).

With the exception of GSE35640^48^ and GSE194040^17^ obtained from microarray, all other datasets employ RNA-seq. For the microarray data, GSE194040^17^ was preprocessed in the original study and downloaded from GEO at the gene level. Only patients who received paclitaxel or the combination of paclitaxel and pembrolizumab were included in this study. Additionally, they were grouped based on whether they were TNBC or not. GSE35640^48^ was downloaded from GEO at the probe set level and then converted to the gene level. For genes with multiple probe sets, we selected the probe set with the highest average hybridization intensity to represent the corresponding gene.

For the RNA-seq data, phs000452.v2.p1^49^, phs000452.v3.p1^50^, phs001493.v1.p1^51^, Gide et al.^52^, and GSE126044 were preprocessed by Lee et al. and requested via ZENODO repository (https://zenodo.org/record/4540874)^53^. Processed RNA-seq gene expression data represented as fragments per kilobase of transcript per million mapped reads (FPKM) of GSE78220^54^ and GSE91061^55^, transcripts per million (TPM) of GSE166449^53^ and counts of GSE165252^56^ were downloaded from GEO. Processed RNA-seq gene expression data from Snyder et al.^57^ was downloaded via ZENODO repository (https://zenodo.org/records/546110). Processed RNA-seq gene expression data represented as TPM were requested from the European Genome-phenome Archive (EGA) with study accession number EGAS00001004343^58^, EGAS00001005013^59^, and EGAS00001004353^60^. For the RNA-seq FASTq data requested from EGAS00001005503^61^, we mapped raw reads to the human genome GRCh38 using HISAT2 (v2.2.1) (https://daehwankimlab.github.io/hisat2/)^62^, with Gencode release 41 as the transcript annotation. Raw read counts for each gene were obtained using featureCounts from the Subread Package (v2.0.1) (https://subread.sourceforge.net/)^63^. The transcripts per kilobase million (TPM) values for each gene were then calculated using an in-house transcript.

These datasets encompassed a variety of cancer types, including skin cutaneous melanoma (SKCM)^48–50,52,55^, non-small cell lung cancer (NSCLC)^53,58,59,64^, renal cell carcinoma (RCC)^51,58,60^, metastatic urothelial carcinoma (mUC)^57,58^, hepatocellular carcinoma (HCC)^61^, breast cancer (BC)^17^, and esophageal cancer (EC)^56^.

Among these datasets, the treatment regimens administered to the patients comprised a wide range of immunotherapy options. This encompassed cancer vaccination (MAGE-A3 vaccine), ICI monotherapy such as anti-CTLA4 (IPI: ipilimumab), anti-PD-1 (NIV: nivolumab, PEM: pembrolizumab), and anti-PD-L1 (ATEZO: atezolizumab), as well as combinations within ICIs. Combination therapies integrating ICIs with chemotherapy, radiotherapy, and targeted therapy such as anti-VEGF (BEV: bevacizumab) were also included. Besides, we incorporated chemotherapy such as docetaxel (TXT) and paclitaxel (PTX) and targeted therapy such as sunitinib (SUN) and sorafenib (SOR), which are treatment arms from the same clinical trials, as control groups in our analysis. The details of treatment information were listed in Supplementary Table 4 (columns “Targets”, “Drug”, “Therapy Type”).

The cohorts from those datasets were grouped according to the cancer type, treatment type, and clinical trials. We displayed the datasets by combining the dataset ID, publication author, or clinical trial name with drug abbreviations. The vast majority of samples, accounting for 97.3%, are pre-treatment samples, denoted either with or without the label “pre”. If the on-treatment and post-treatment samples possess a sufficient cohort size for analysis, they will be separate datasets with the labels “on” or “post”. The detailed sample information was described in the Supplementary Table 4 (columns “Platform”, “Sample Type” and “Boipsy”).

To define the responders and non-responders, our objective was to adhere closely to the criteria outlined in the original studies. In cases where response criteria were defined within specific clinical trial cohorts, we adhered to the original definitions provided (phs000452.v2.p1, GSE35640, GSE166449, GSE126044, GSE165252). The other cohorts with available RECIST information, we classified complete response (CR) and partial response (PR) as responders, and stable disease (SD) and progressive disease (PD) as non-responders. For the dataset phs000452.v3.p1, we designated CR, PR, and MR as responders, and SD and PD as non-responders. In the case of GSE194040, we categorized pathological complete response (pCR) as responder and non-pCR as non-responder. Patients lacking clear response information were excluded from the analysis. The definition of each dataset (columns “Def. Responder(1)” and “Def. Non-responder(0)”) and response distribution (columns “No. Samples”, “No. Non-responder(0)”, “No. Responder(1)”) were also described in the Supplementary Table 4.

### Framework of TimiGP-Response

As described before, we presented TimiGP framework to analyze the association between Tumor Immune Microenvironment (TIME) and prognosis, which was designed to infer the cell-cell interaction networks and prioritize the cell type associated with prognosis (favorability score) (TimiGP-Prognosis)^14,15^.

In brief, TimiGP-Prognosis required three inputs: (1) immune cell-type markers, which can be obtained from different sources, e.g., prior knowledge, bulk transcriptome clustering or single cell RNA-seq; (2) transcriptomic profiles, e.g., bulk RNA-seq or microarray; and (3) survival statistics from the same cohort, including event (e.g., death, recurrence) and time-to-event. The algorithm of TimiGP-Response involves five steps: Step 1 - Define and select marker gene pair matrix; Step 2 - choose gene pairs associated with favorable prognosis; Step 3 - Construct the directed gene-gene network; Step 4 - Identify cell-cell interactions by enrichment analysis; and Step 5 - Build and analyze the cell-cell interaction network. More details are available in our previous publication^14,15^.

According to the similar rationale that the dynamic balance of pre-existing immunity will affect patient’s response to immunotherapy (Supplementary Fig. 1A), we tailored the TimiGP framework originally proposed in prognosis (TimiGP-Prognosis)^14,15^ to therapy response (TimiGP-Response) through the following adjustments (Supplementary Fig. 1B):

1) Input - Response information: The third input, originally based on survival statistics, was adjusted to reflect immunotherapy response, a binary variable denoted by 0 for non-response and 1 for response.
2) Step 2 - choose gene pairs associated with responders: The second step of the algorithm, which involved using Cox regression to select marker pairs associated with favorable prognosis (‘TimiCOX’ Function), was replaced by Fisher’s exact tests (‘TimiFisher’ Function). This change was made to identify marker pairs significantly associated with responders, considering the binary nature of both marker pair scores and clinical information, as well as the limited proportion of responders.

All other steps in the framework remain unchanged, and their details are available in our previous publication on TimiGP-Prognosis^15^. To implement the framework, TimiGP-Response is included in version 1.3.0 of the TimiGP R package, which was upgraded in this study. It is available on GitHub at https://github.com/CSkylarL/TimiGP.

Correspondingly, the two outputs of TimiGP-Response now signify the correlation between the TIME and therapy response, rather than a favorable prognosis (Supplementary Fig. 1B). Within the cell-cell interaction network, for instance, the notation X → Y denotes a higher cell X/Y ratio or increased functionality linked with responders. A higher favorable score signifies that the respective cell type is typically associated with responses, whereas a higher unfavorable score suggests the cell type is associated with non-responses.

### Customized TimiGP-Response analysis with scRNA-seq in TNBC

#### Identification of cell types from scRNA-seq

The single-cell RNA-seq data (GSE169246) was preprocessed in the original study^16^ and was loaded and analyzed by Seurat R Package (v4.3.0)^65^. For general quality control, the preprocessed data passed the filter to retain cells with a number of detected genes between 200 and 8000 and a mitochondrial gene percentage of 10% or less. The RNA expression was log normalized with a scale factor of 100,000 (‘NormalizeData Function’). Due to poor data quality compared to others, we excluded patients P010 and P016. Given the limited number of pre-treatment chemo-immunotherapy samples and our goal to obtain consistent cell-type markers that adhere to the IMC data^18^, we analyzed the remaining datasets, which included both treatments and pre- and post-treatment samples.

We identified 2,000 highly variable features using the ‘vst’ selection method (‘FindVariableFeatures Function’), and then scaled the data, regressing out the effects of RNA count and mitochondrial percentage (‘ScaleData Function’).

For clustering, we performed Principal Component Analysis (PCA) for dimension reduction(‘RunPCA Function’) and selected the first 30 PCs to construct a Shared Nearest Neighbor (SNN) graph with a k parameter of 20 (‘FindNeighbors Function’). Clustering was defined at a resolution of 0.8. The major immune cell types (“CD20+B”, “CD79a+Plasma”, “cDC”, “pDC”, “M1”, “M2”, “Mast” and “NK & T”) were defined based on their marker expression (Fig. 1e and Supplementary Fig. 3).

The cells from the “NK & T” cell cluster were extracted to define the subtypes. Similar to the previous procedure, the subsetted data was then clustered using the first 30 PCs and a k parameter of 20 at a resolution of 1. The NK cells (“CD56+NK”) and T cell subtypes (“CD4+TCF1+T”, “CD4+PD1+T”, “Treg”, “CD8+TCF1+T”, “CD8+GZMB+T”, CD8+PD1+Tex”) were defined based on their marker expression (Fig. 1e and Supplementary Fig. 3). Uncharacterized clusters were dropped for TimiGP-Response analysis.

For visualization, we applied the Uniform Manifold Approximation and Projection for Dimension Reduction (UMAP), using the first 30 PCs, a minimum distance of 0.8, and 30 neighbors, with a set seed of 1234 to ensure reproducibility (‘RunUMAP’ Function).

#### Customization of cell-type markers from scRNA-seq

After defining the cell types, we identified cell-type-specific markers using the ‘FindAllMarkers’ Function from the Seurat R package with the preprocessed scRNA-seq data mentioned earlier. Multiple methods were employed to choose approximately 30 of the most specific markers for each cell type.

First, we utilized the Receiver Operating Characteristic (ROC) method. Each gene is assessed individually by constructing a classifier based solely on that gene to discriminate between two groups of cells (the cell type itself and all others), using the Area Under the Curve (AUC) as the evaluation metric. We set the criterion of an AUC > 0.7 for cell types including “Treg”, “CD8+GZMB+T”, “CD8+PD1+Tex”, “CD56+NK”, and “CD20+B”. However, for “cDC”, “pDC”, “M1”, “M2”, “Mast”, and “CD79a+Plasma”, AUC > 0.7 was considered too weak, so we chose the top 30 markers based on AUC.

Second, we employed the Wilcoxon rank-sum test to detect genes that are differentially expressed between each cell type and all others. For cell types including “CD4+TCF1+T”, “CD8+TCF1+T”, and “CD4+PD1+T”, where too few markers were available from the ROC method, we appended the top 30 markers based on p-value and average log2 fold change in the Wilcoxon results.

Finally, we manually included markers utilized to define these cell types from scRNA-seq data and the corresponding genes for IMC protein markers^18^, though the majority of them had been incorporated in the preceding two steps.

Following these steps, we generated the final set of markers customized for TimiGP-Response analysis.

#### Customized TimiGP-Response analysis in TNBC

The TimiGP-Response analysis was performed with the TimiGP R package (v1.3.0) upgraded in this study. The above cell-type annotation customized for the TNBC and anti-PD-1/PD-L1 treatment was then used as the first input for TimiGP-Response to analyze the second input, TNBC cohort treated with a combination of chemo and anti-PD-1 therapy from the microarray data (GSE194040^17^), along with the corresponding treatment response, the third input. Using ‘TimiFisher’ and ‘TimiEnrich’ Functions, we selected 4480 (∼top 10% following p-value) immune marker gene pairs (IMGPs) significantly associated with responders (P-Value < 0.05) and defined the cell-cell interactions with default parameters (false discovery rate [FDR] < 0.05, where the FDR is calculated by adjusting the p-values from the enrichment analysis using the Benjamini-Hochberg method). Fig. 1c was generated using ‘TimiCellChord’ Function; The favorability score in Fig.1d was calculated with ‘TimiFS’ Function and visualized using ‘TimiFSBar’ Function.

### Comparison of TimiGP and IMC Results in TNBC

#### Evaluation of immune cell markers: scRNA-seq vs. IMC

The IMC data(https://doi.org/10.5281/zenodo.7990870) was processed to obtain cell-level information and corresponding panels according to the original study’s methodology^18^. We adhered to the established thresholds for positive markers and the annotations for cell clusters. The protein expression values were scaled and clipped at the 99th percentile to mitigate the impact of outliers. For each cell type and protein, the median of these scaled values was then calculated to provide a robust measure of central tendency. These medians underwent additional z-score normalization, individually applied to each protein within every cell type.

We narrowed our focus to 11 immune cell types, specifically “CD20+B”, “CD79a+Plasma”, “CD4+TCF1+T”, “CD4+PD1+T”, “Treg”, “CD8+TCF1+T”, “CD8+GZMB+T”, “CD8+PD1+Tex”, “CD56+NK”, “cDC”, and “M2”, while excluding non-relevant markers. We then visualized the normalized protein expression levels in a heatmap (Fig. 1e).

The expression levels of genes corresponding to the IMC protein markers from the scRNA-seq data (GSE169246^16^) were scaled and visualized in a dot plot. This dotplot, created using the ‘DotPlot’ Function from the Seurat R package, depicts both the gene expression percentage and average expression for each cell type (Fig. 1e). Cell types defined by both IMC and scRNA-seq consistently exhibit similar expression patterns, facilitating comparisons between IMC and TimiGP results.

#### Analysis of cell-response associations and cell-cell interactions using IMC data

After confirming the consistent definition of 11 immune cell types between scRNA-seq and IMC data, we further analyzed the IMC data to orthogonally validate TimiGP results. We retrieved the cell count and density for each cell type per patient from the preprocessed IMC data. Samples were grouped by treatment and biopsy time point and the logistic regression was performed within each group to assess the association of immune cell density and cell ratio with response.

For each cell density, the logistic regression was performed using the ‘glm’ Function from the glmnet R package (v4.1-6)^66^ with ‘family = “binomial“’ for binary outcome analysis. An odds ratio significantly greater than 1 indicates a positive association, while an odds ratio less than 1 suggests a negative association. Significance was defined as FDR < 0.05, where FDR represents p-values adjusted using the Benjamini-Hochberg (BH) method to account for multiple comparisons.

Given that TimiGP interactions partially represent associations between cell ratios and responders, we also computed the log2 ratios between each pair of cell types within each group using cell count. For each cell ratio, we fitted a logistic regression model to evaluate its association with response information in a similar way. The cell-cell interactions were defined with Odds ratios > 1 and FDR < 0.1.

#### Comparison between TimiGP and IMC results

Following the re-analysis of the IMC data, a comparative assessment was conducted between the IMC findings and those inferred through TimiGP. To compare the contribution of cell types to treatment response, associations between 11 cell types and response from IMC data, were visualized using a heatmap generated with the ComplexHeatmap (v2.15.1) R package (Supplementary Fig. 5). In pre-treatment/baseline samples from both methodologies and cohorts, CD8+ GZMB+ T cells emerged as notably associated with responders.

Given the inherent limitations in statistical power stemming from IMC data quality and technological constraints, we further compared cell-cell interactions that represent a high ratio between two cell types associated with responders. The 16 TimiGP cell-cell interactions that were orthogonally validated by IMC findings were used to hierarchical network using Cytoscape (v3.10.1)^67^ (Fig. 1f). Statistical attributes of these interactions, encompassing enrichment analysis for TimiGP cell-cell interactions and logistic regression for IMC cell-cell interactions, were visualized through a dot plot utilizing the ggplot2 R package (v3.4.1)^68^ (Fig. 1g).

### Pan-cancer TimiGP-Response analysis

#### Multi-resolutions analysis with different cell-type annotations

For the bulk transcriptomics datasets collected for pan-cancer analysis (Supplementary Table 4), we applied TimiGP-Response to each dataset separately through the TimiGP R package (v1.3.0) upgraded in this study. We conducted the pan-cancer analysis at different resolutions and for various purposes using modified cell-type annotations from Bindea2013 (immunoscore)^19^, Newman2015(LM22)^20^, Zheng2021(pan-cancer T cell scRNA-seq)^30^, all of which are incorporated into the TimiGP R package^14,15^. Bindea2013, which includes positive controls - anti-tumor cytotoxic cells and negative control associated with non-responders - tumor cells, was used to examine the feasibility of TimiGP-Response in pan-cancer analysis. Newman2015 was employed to profile the pan-cancer landscape at the tumor immune microenvironment (TIME) level, including innate and adaptive immune cell types and states. Zheng2021 was utilized to portray the pan-cancer association of T cell subtypes with therapy response.

#### Automatic determination of responder-associated IMGPs in Step 2 of TimiGP-Response

Considering the varying quality and characteristics between datasets, we implemented a series of condition-based cutoffs tailored to each dataset to determine the immune marker gene pairs (IMGPs) associated with responders. Initially, we applied the default cutoff FDR < 0.05. If the proportion of selected IMGPs is less than 1%, a more lenient condition p-value < 0.01 will be used as a cutoff. If the proportion of selected IMGPs is still less than 1%, an even more permissive condition p-value < 0.05 will be employed. For specific datasets including “Liu_phs000452.v3.p1”, “Wolf_GSE194040_ISPY2_ptx_pem_non-TNBC”, “Wolf_GSE194040_ISPY2_ptx_pem_TNBC”, and “Wolf_GSE194040_ISPY2_paclitaxel_TNBC”, we manually set a cutoff of PV < 0.05. For “Wolf_GSE194040_ISPY2_paclitaxel_non-TNBC”, a stricter cutoff of FDR < 0.01 was utilized. This approach automatically balances the statistical power between this step and subsequent analyses.

#### Definition of cell-cell interactions

All datasets primarily utilize the default FDR < 0.05 threshold to ascertain cell-cell interactions. However, in instances where datasets exhibit suboptimal data quality, such as a limited number of samples leading to insufficient cell-cell interactions for analysis, we manually determined the cutoff for cell-cell interactions.

For “GSE165252on” and “GSE165252post”, analyzed with the Bindea2013 cell-type annotation, cell-cell interactions were identified with a threshold of P-value < 0.05. For “GSE91061pre: NIV/PEM” and “GSE166449: PEM”, analyzed with the Zheng2021 cell-type annotation, cell-cell interactions were identified with a threshold of P-value < 0.01. For “GSE165252post”, analyzed with both the Zheng2021 and Newman2015 cell-type annotations, cell-cell interactions were assessed using gene sets with thresholds of P-value < 0.05 and P-value < 0.01, respectively.

#### Integration of pan-cancer results

For every dataset, the cell-cell interaction network was constructed using the ‘TimiCellChord’ function, and the favorability score was computed using the ‘TimiFS’ function and presented visually through the ‘TimiFSBar’ function. The detailed chord diagram illustrating the cell-cell interaction network and the barplot displaying the favorability score for each dataset are conveniently accessible in a database-like format via the following link: ahttps://github.com/CSkylarL/MSofTimiGP-Response. To integrate the pan-cancer results, we visualized the favorability score by scatterpie R package (v0.1.8)^69^ (Fig. 2).

### Calculation of similarity between cell-cell interaction networks

The similarity between these networks (e.g., between A and B) was calculated by according to Tversky index:

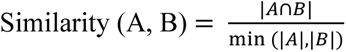

In the equation, |*A* ∩ *B*| is the number of shared interactions (e.g., Cell X → Cell Y) between network A and B, and the min (|*A*|, |*B*|) is the largest number of interactions that can be shared by A and B. The index takes a value within [0,1] with 0 and 1 indicating the lowest and the highest similarity, respectively.

The statistical significance of the similarity is evaluated based on the hypergeometric distribution:

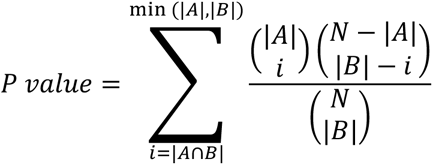

Where N is the total number of potential cell-cell interactions annotated by given cell-type markers.

### Quantification and statistical analysis

The major analysis were based on R (v4.2.0) and R libraries (dplyr (v1.1.0)^70^, reshape (v0.8.9)^71^, data.table (v1.14.8)^72^, doParallel (v1.0.17)^73^, foreach (v1.5.2)^74^, scales (v1.2.1)^75^, ggplot2 (v3.4.1)^68^, scatterpie (v0.1.8)^69^, gridExtra (v2.3)^76^, glmnet(v4.1-6)^66^, ComplexHeatmap (v2.15.1)^77^, RColorBrewer (v1.1-3)^78^, circlize (v0.4.15)^79^, Seurat(v4.3.0)^65^, Tibble(v3.1.8)^80^, Matrix(v 1.5-3)^81^, ggpubr(v 0.6.0)^82^, fst(v 0.9.8)^83^. The TimiGP R package (v1.3.0) was upgraded in this study with the help of roxygen2 (v7.1.1)^84^ and devtools (v2.4.2)^85^ R packages. The hierarchical network in Fig. 1f was visualized by Cytoscape (v3.10.1)^67^.

### Data and code availability

The preprocessed IMC data were requested from via Zenodo data repository (https://doi.org/10.5281/zenodo.7990870). The preprocessed scRNA-seq data was obtained from GEO with accession number GSE169246 (https://www.ncbi.nlm.nih.gov/geo/query/acc.cgi?acc=GSE169246). All bulk RNA-seq and microarray data are publicly available. Accession numbers for these datasets are listed in Supplementary Table S4.

The TimiGP R package (v1.3.0) upgraded in this study is publicly available as of the publication date for non-profit academic use (https://github.com/CSkylarL/TimiGP). The cell type markers used in this study were integrated into the R package.

All code used in this study, along with detailed TimiGP-Response results for each dataset, are available on GitHub: https://github.com/CSkylarL/MSofTimiGP-Response. To facilitate replication of the analysis code, some preprocessed and intermediate data have been deposited in the Zenodo data repository. The analyzed scRNA-seq data (GSE169246), including cell type information, is publicly available (https://doi.org/10.5281/zenodo.12209783). Additionally, example bulk transcriptomics datasets for TimiGP-Response can be accessed at https://doi.org/10.5281/zenodo.12209773. Any additional information required for this study is available upon request.

## Acknowledgments

This work was supported by the National Cancer Institute of the National Institute of Health Research Project Grant (J.Z., R01CA234629-01), the AACR-Johnson & Johnson Lung Cancer Innovation Science Grant (J.Z., 18-90-52-ZHAN), the MD Anderson Physician Scientist Program(J.Z.), the MD Anderson Lung Cancer Moon Shot Program(J.Z.) and the Cancer Prevention Research Institute of Texas (CPRIT) (C.C., RR180061). C.C. is a CPRIT Scholar in Cancer Research. A.M. is supported by the Sheikh Khalifa Bin Zayed Al-Nahyan Foundation and MD Anderson Pancreatic Cancer Moonshot. The authors would like to acknowledge the support of the High Performance Computing for research facility at the University of Texas MD Anderson Cancer Center for providing computational resources that have contributed to the research results reported in this paper. The Fig.1a, Fig. 2a, Supplementary Fig. 1 were created with BioRender.com. We would like to express special thanks to Dr. Baoyi Zhang and other previous lab members who help downloaded some datasets when they studied in Dr. Chao Cheng’s lab.

## Author information

### Contributions

C.L. conceptualized this study and developed analysis strategies. C.L. developed the R package and performed the formal analysis. C.L., and C.C. assisted with data acquisition. W.H preprocessed RNA-seq FASTq data requested from EGAS00001005503 to generate TPM gene expression. A.M., L.W., A.R., J.Z. and C.C. provided expertise and feedback. C.L. wrote the original draft. J.Z. and C.C. reviewed and edited the manuscript. All authors commented on the manuscript at all stages. C.C. and J.Z. co-supervised this study and provided funding resources.

