## Supplementary Figures for "TimiGP-Response: the pan-cancer immune landscape associated with response to immunotherapy"

**a** The TimiGP-Reponse Rationale

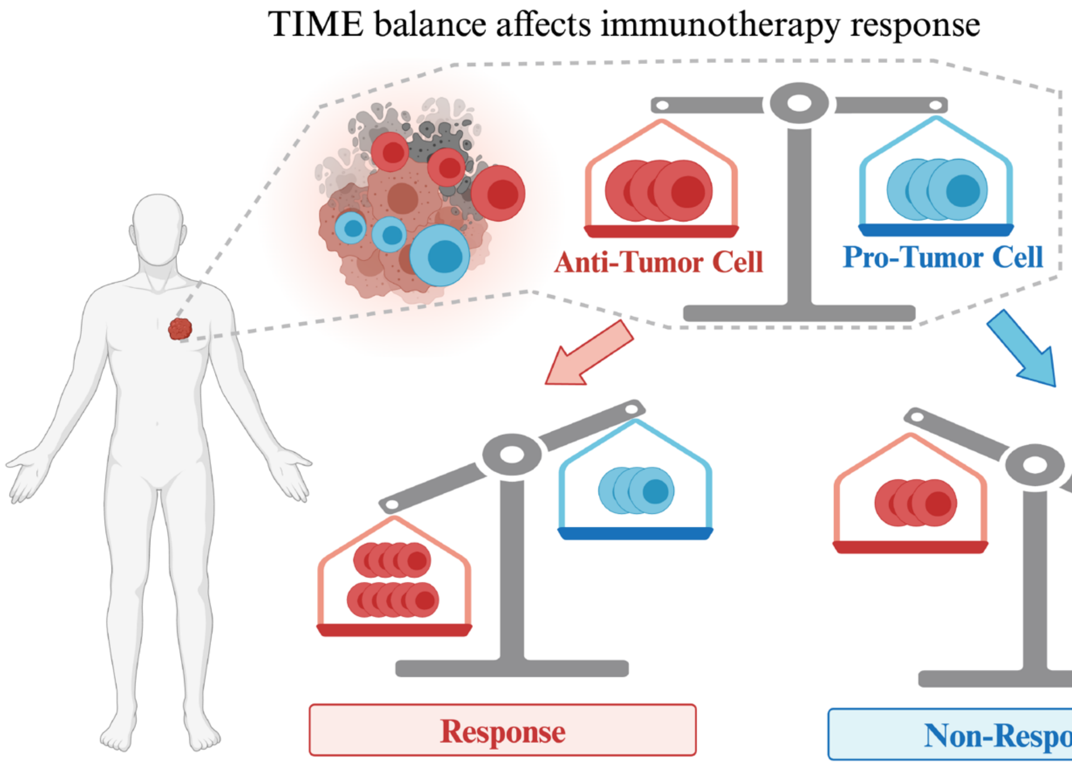


**b** The TimiGP-Reponse Framework

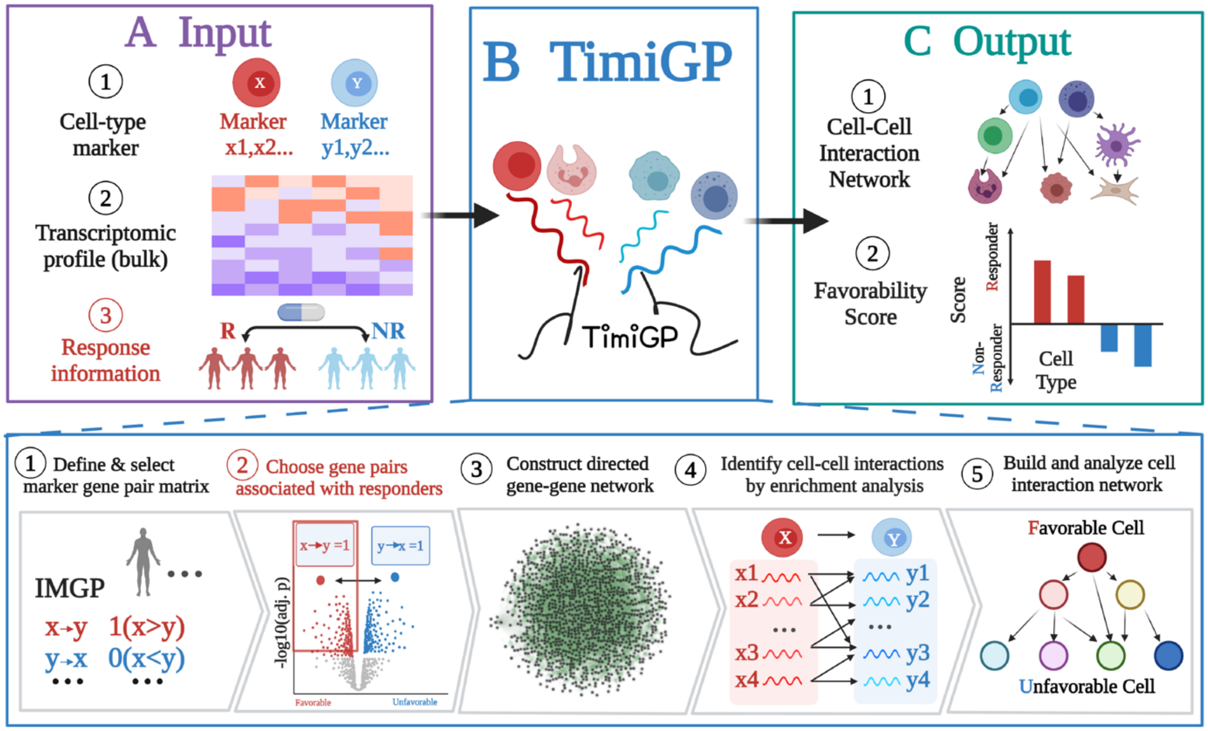

**Supplementary Figure 1 | The rationale and framework of TimiGP-Response adapted from TimiGP-Prognosis. Related to Figure 1**

(a) Representative schema of TimiGP - Response Rationale. The tumor immune microenvironment (TIME) is a balance between anti-tumor and pro-tumor immune cells. Such pre-existing immunity will affect patient's response to immunotherapy. If the function of anti-tumor cell types is more vital than the pro-tumor cells (e.g., higher abundance, higher marker expression), the TIME is associated with responders; otherwise, it is associated with unfavorable non-responders. This figure was created with BioRender.com.

(b) Schematic depicting the entire framework of the TimiGP - Response. The red text highlights the framework modifications: 1) The third input is response information, and 2) the second step involves selecting gene pairs associated with responders. Correspondingly, the generated cell-cell interaction network is associated with the responder, and the calculated favorable score indicates response, while the unfavorable score indicates non-response. This figure was created with BioRender.com.

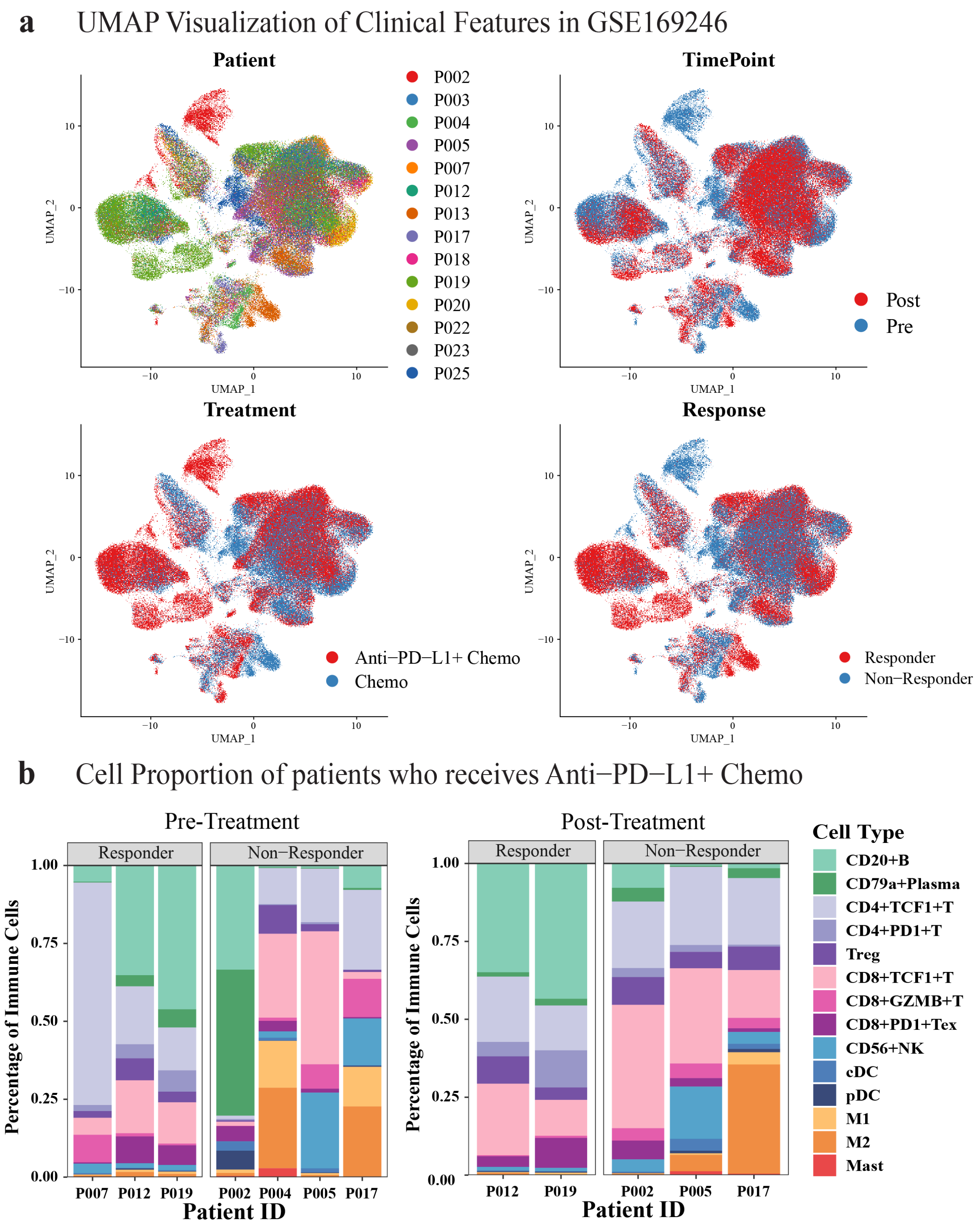


**Supplementary Figure 2 | Overview of clinical features and cell proportions reanalyzed from GSE169246. Related to Figure 1**

(a) UMAP visualization of clinical features of single-cell RNA-seq data for TNBC (GSE169246). (b) Cell proportion of patients who receive the combination of anti-PD-L1 treatment and chemotherapy (Left: pre-treatment samples; Right: post-treatment samples).

UMAP of cell-type marker expression

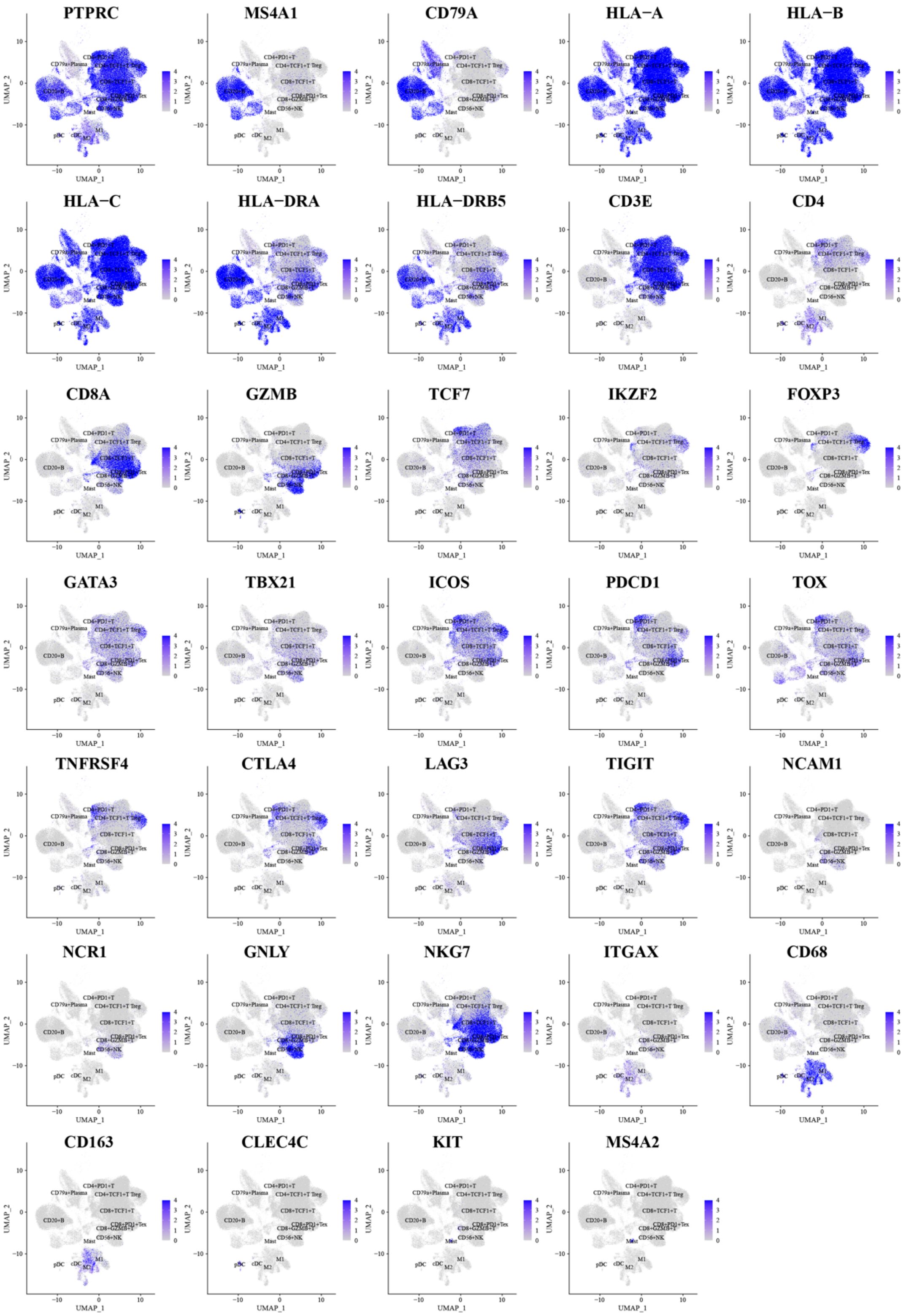


**Supplementary Figure 3 | The expression of cell-type markers in UMAP. Related to Figure 1**

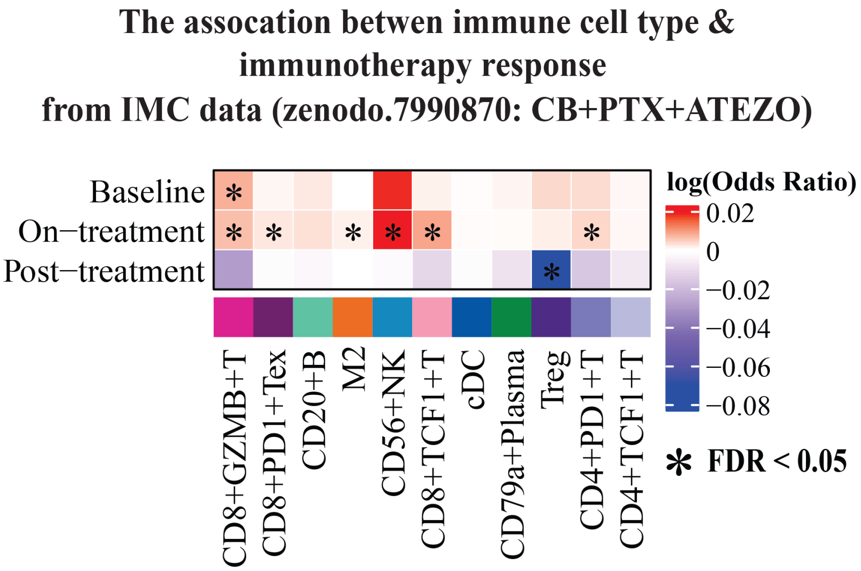


**Supplementary Figure 4 | The association between immune cell density and immunotherapy response**

The relationship between immune cell density and immunotherapy response was assessed using univariate logistic regression across different biopsy time points in the IMC dataset (zenodo.7990870: CB+PTX+ATEZO). The color scale indicates the log-transformed odds ratio: values greater than 0 indicate a positive association, whereas values less than 0 indicate a negative association. The significance was defined as FDR < 0.05, where FDR represents p-values adjusted using the Benjamini-Hochberg (BH). Abbreviation of cell types: B, B cell; Plasma, Plasma cell; T, T cell; Treg, Regulatory T cell; Tex, Exhausted T cell; NK, Natural killer cell; cDC, Conventional Dendritic cell; M2, Anti-inflammatory macrophages.

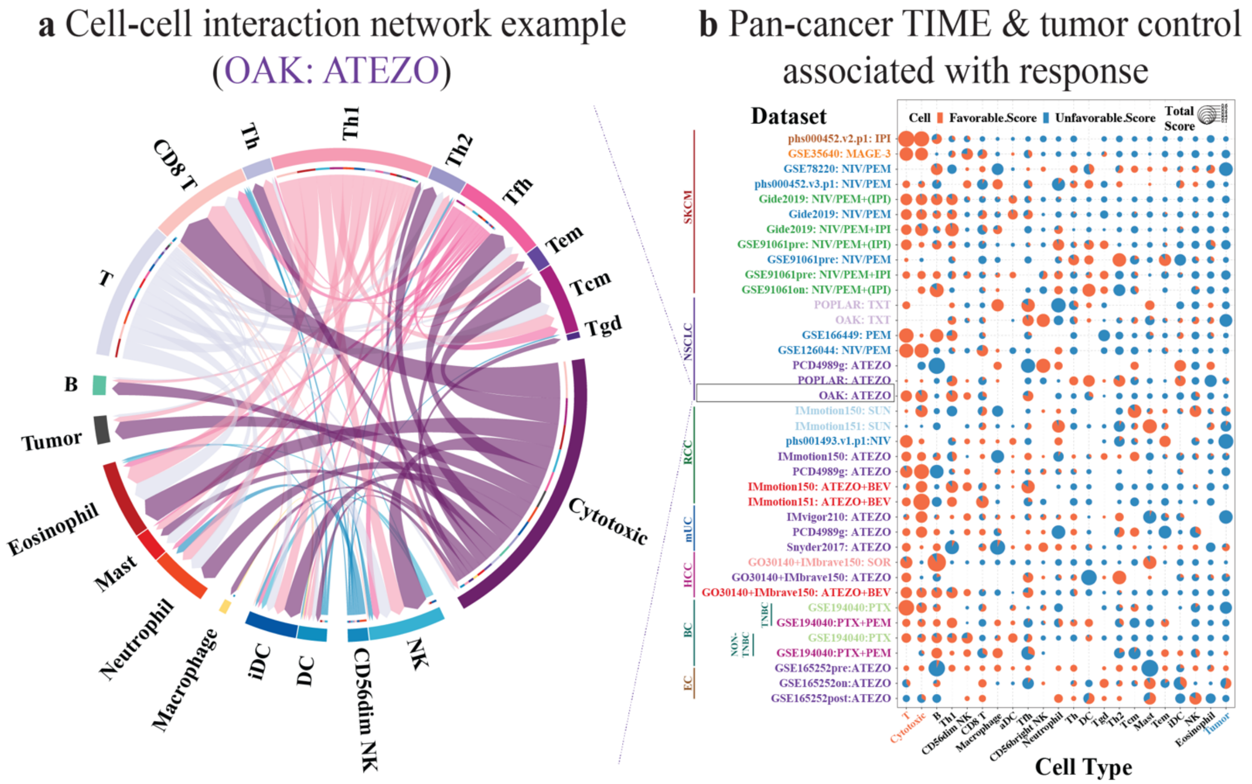


**Supplementary Figure 5 | Robust performance of TimiGP-Response in pan-cancer analysis. Related to Figure 2**

(a) An example of cell-cell interaction network. The dataset is OAK:ATEZO from panel b as indicated with dashed lines. (b) The scatter pie chart displays the favorability score exported by TimiGP - Response Module, which estimates the association of immune cell types and tumor cells defined by modified Bindea2013 markers (x-axis) with immunotherapy responders (favorable score, orange) and non-responders (unfavorable score, blue). The cell types discussed in results were highlighted with orange and blue colors. There are 7 cancer types in this analysis: SKCM (melanoma), NSCLC (non-small cell lung cancer), RCC(renal cell carcinoma), mUC (metastatic urothelial cancer), HCC (hepatocellular carcinoma), BC (breast cancer; TNBC, triple-negative breast cancer), EC (esophageal cancer). The datasets (y-axis) are labeled in the format "dataset ID: drug", with immunotherapy datasets shown in dark colors, chemotherapy and targeted therapy datasets shown in light colors. The immunotherapy includes anti-CTLA4 (IPI: ipilimumab), MAGE-A3 vaccine, anti-PD-1 (NIV: nivolumab, PEM: pembrolizumab), combination of anti-CTLA4 and anti-PD-1, anti-PD-L1 (ATEZO: atezolizumab), and combination of anti-PD-L1 and anti-VEGF (BEV: bevacizumab). The chemotherapy includes docetaxel (TXT) and paclitaxel (PTX). The targeted therapy includes sunitinib (SUN) and sorafenib (SOR) which target receptor tyrosine kinases (RTKs).

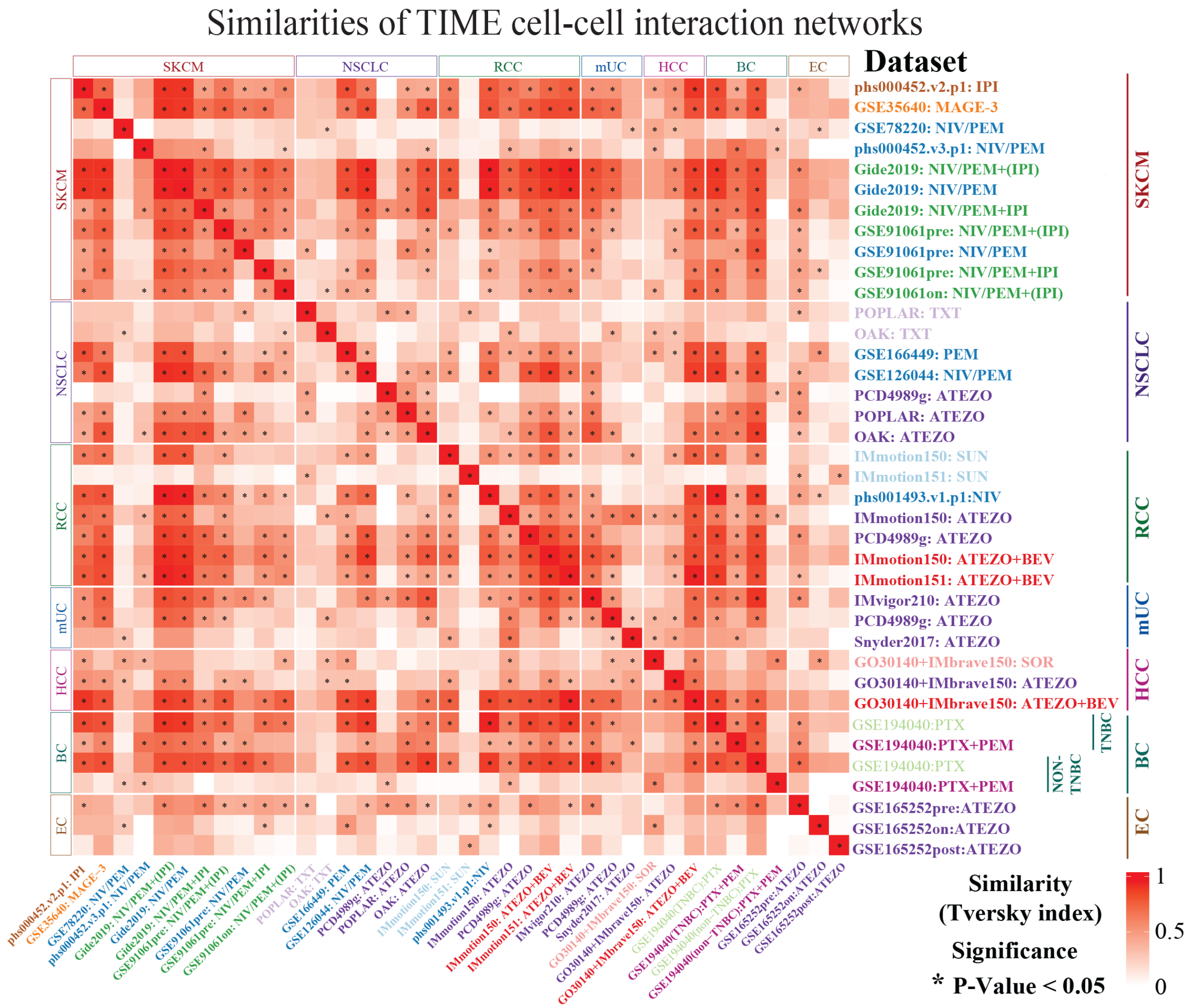


**Supplementary Figure 6 | High similarities of TIME cell-cell interaction networks across cancer types and treatments. Related to Figure 2**

Cell-cell interactions were exported for each cohort using TimiGP-Response at TIME level with LM22 signature. Consistency among these networks was determined through Tversky index calculation, while statistical significance of similarity was assessed using the hypergeometric distribution.
